# Contribution of Implicit Memory to Adaptation of Movement Extent During Reaching Against Unpredictable Spring-Like Loads: Insensitivity to Intentional Suppression of Kinematic Performance

**DOI:** 10.1101/2023.03.13.532441

**Authors:** Devon D Lantagne, Leigh Ann Mrotek, James B Hoelzle, Danny G Thomas, Robert A Scheidt

## Abstract

We examined the extent to which intentionally underperforming a goal-directed reaching task impacts how memories of recent performance contribute to sensorimotor adaptation. Healthy human subjects performed computerized cognition testing and an assessment of sensorimotor adaptation wherein they grasped the handle of a horizontal planar robot while making goal-directed out-and-back reaching movements. The robot exerted forces that resisted hand motion with a spring-like load that changed unpredictably between movements. The robotic test assessed how implicit and explicit memories of sensorimotor performance contribute to the compensation for the unpredictable changes in the hand-held load. After each movement, subjects were to recall and report peak movement extent from the previous trial. Subjects performed the tests under two counterbalanced conditions: one where they performed with their best effort, and one where they intentionally suppressed performance. Results from the computerized cognition tests confirmed that subjects understood and complied with task instructions. When suppressing performance during the robotic assessment, subjects demonstrated marked changes in reach precision, time to capture the target, and reaction time. We fit a set of limited memory models to the data to identify how subjects used implicit and explicit memories of recent performance to compensate for the changing loads. In both sessions, subjects used implicit, but not explicit, memories from the most recent trial to adapt reaches to unpredictable spring-like loads. Subjects did not “give up” on large errors, nor did they discount small errors deemed “good enough.” Although subjects clearly suppressed kinematic performance (response timing, movement variability, and self-reporting of reach error), the relative contributions of sensorimotor memories to trial-by-trial variations in task performance did not differ significantly between the two testing conditions. We conclude that intentional performance suppression had minimal impact on how implicit sensorimotor memories contribute to adaptation of unpredictable mechanical loads applied to the hand.

## Introduction

The human brain utilizes explicit and implicit processes to shape repeated performance of actions such as reaching and pointing (Shadmehr et al. 2010; Krakauer and Mazzoni 2011; Taylor et al. 2014; McDougle et al. 2015). We use the term *explicit* to describe situations where stimuli are consciously perceived and the performer has declarative access to the way the stimuli affect their behavior (c.f., Reber 1989; Reber A.S. et al. 1999; Maresch et al. 2021). We use the term *implicit* when stimuli are not consciously perceived and the performer is unaware how the stimuli impacts behavior. The explicit learning process relies on memories that develop quickly and contribute to large reductions in reach error during adaptation via perceived performance errors (Smith et al. 2006; Kiesler and Shadmehr, 2010; Taylor et al. 2014; McDougle et al. 2015), especially if individuals are aware of the causal source of the error (Werner and Bock 2007; Hegele and Heuer 2010). Guided by conscious decisions, explicit learning can be verbally articulated (Magill 2011), a fact that has been exploited to probe the developmental time course of explicit motor plans and strategies (Taylor et al. 2014; McDougle et al. 2015). Explicit learning is associated with brain activity in the prefrontal, parietal cortex, and hippocampal regions (Scoville and Milner 1957; Coull and Nobre 2008; Ikkai and Curtis 2011; Eriksson et al. 2015; Wolpe et al. 2020). By contrast, the implicit learning process relies on memories that develop slower than those serving explicit processes but can be retained longer. Implicit learning is driven by sensory prediction errors – a discrepancy between sensory feedback and the brain’s prediction of expected feedback (Mazzoni and Krakauer 2006; Hwang et al. 2006; McDougle et al. 2015). This process is automatic, ongoing, and operates unconsciously in a way that is inaccessible to verbal description (Frensch 1998; Magescas and Prablanc 2006). Implicit learning mechanisms have been associated with brain activity throughout posterior parietal cortices, cerebellum, and basal ganglia (Diedrichsen et al. 2005; Scheidt et al. 2012; Izawa et al. 2012; Reber, 2013).

Explicit and implicit learning appear to operate simultaneously (Smith et al. 2006; Taylor et al. 2014; Bond and Taylor 2015; McDougle et al. 2015) and interact in complex ways. Careful experiments have shown that implicit learning can interfere with the efficacy of explicit strategies (Mazzoni and Krakauer 2006). Likewise, employing an explicit strategy to counteract an environmental perturbation can impede the development of implicit learning (Benson et al. 2011), although it is not yet known whether this effect may be caused by a reduction in the magnitude of errors driving implicit learning or by inhibition of the implicit learning processes themselves. Nevertheless, it is evident from these past works that some experimental tasks appear better able to cause interference between explicit and implicit learning mechanisms than others. In most previous studies, explicit and implicit learning have been assayed using measures of movement errors induced by deterministic perturbations (e.g., predictable visuomotor rotations, prism goggles, static force fields). In the present work, we sought to isolate implicit sensorimotor learning using unpredictable but biased perturbations to discourage engagement of explicit learning mechanisms (cf., Scheidt et al. 2001; Judkins and Scheidt 2014; Lantagne et al. 2021). We previously showed that under such conditions, adaptation of goal-directed reach extent relies on implicit memories of kinematic error to the exclusion of explicitly recalled errors (Lantagne et al. 2021). Using this approach, we now seek to assay potential interactions between explicit and implicit learning processes by determining the extent to which implicit adaptation is sensitive to an explicit strategy of intentional suppression of kinematic performance.

More specifically, we tested the hypothesis that contributions of implicit memories to sensorimotor adaptation can be resistant to explicit interference through intentional performance suppression. Healthy adult subjects with no known history of neuromotor injury grasped the handle of a horizontal planar robot while performing repeated goal-directed reaches under two task conditions: with best effort, and while attempting to suppress performance by mimicking mild traumatic brain injury according to instructions adapted from Suhr and Gunstad (2000). The robot opposed the reaches with spring-like loads that changed in strength between each reach in an unpredictable manner. This seemingly random aspect of the experimental task encourages the preferential engagement of implicit learning mechanisms and discourages the use of an explicit strategy (Lantagne et al. 2021). After each reach, we asked subjects to recall and report the actual extent of their movement on the most recent attempt. We analyzed movement kinematics and the ability of several memory-based models of sensorimotor adaptation to account for variations within the kinematic data (c.f., Scheidt et al. 2001, 2012; Scheidt and Stoeckmann 2007; Judkins and Scheidt 2014; Lantagne et al. 2021). These models assessed the possibility that explicit and/or implicit memories of prior performance impacted subsequent reaches to differing degrees under the two task conditions. They also considered the possibility that reach adaptation might involve discounting of large or small errors (c.f., Goodrich et al. 1998; Körding and Wolpert 2004). We assessed the extent to which intentional performance suppression impacted kinematic performance and the extent to which it impacted how implicit and explicit sensorimotor memories contribute to trial-by-trial corrections for reach performance errors. Our findings support the conclusions that the adaptation of reach extent is driven primarily by implicit memories of sensorimotor performance when environmental loads change randomly from one trial to the next, and that despite marked degradation of reach performance during conscious attempts to suppress performance, the way implicit sensorimotor memories contribute to adaptation remains unchanged.

## Methods

### Participants

Nineteen healthy young adults provided written, informed consent to participate in this study. The cohort comprised 8 females and 11 males [age: 24.3 ± 0.6 years (mean ± 1 standard deviation, here and elsewhere)]. All had normal or corrected-to-normal vision and none had any known neurological deficits. Subjects were recruited from the Marquette University student population. All experimental procedures received institutional review and approval in accordance with the Declaration of Helsinki (protocol HR-3233).

### Experimental Setup and Procedures

All subjects participated in two experimental sessions lasting about 45 minutes each. The sessions were typically performed on the same day but were separated in time by at least 1 hour. In one session, subjects were to perform the experimental tests to the best of their ability (the Best Effort condition; BE). In the other session (the Performance Suppression condition; PS), we adapted an approach established by Suhr and Gunstad (2000) whereby subjects were instructed to suppress performance by emulating symptoms typically associated with concussion (see Appendix A). These instructions described eight common symptoms of concussion including: slowed responses; difficulty concentrating; memory problems; fatigue; headache; irritability; anxiety; and depression. After testing was completed for the Performance Suppression session, subjects were asked to rate the extent to which they attempted to emulate each symptom using a Likert-like scale that ranged from 0 to 10, with lower values indicating less effort. Condition order (BE or PS) was counter-balanced across subjects.

Within each session, subjects performed two experimental tests. The first was a subset of tests drawn from a clinically accepted computerized tool designed to assess several aspects of cognitive performance (CogState, Inc, Melbourne, Australia). The purpose of the cognition tests was to verify that subjects understood and complied with task instructions in the two sessions and to identify how performance suppression would manifest in the speed and accuracy of test performance. The second test was an established robotic assessment of sensorimotor adaptation during goal-directed reaching (Lantagne et al. 2021). The purpose of this test was to identify how performance suppression would manifest in goal-directed reaching against unpredictable mechanical loads, and to determine whether the suppression of performance would impact memory mechanisms supporting adaptation to changing loads.

#### Computerized Cognitive Assessment

Subjects were seated in a quiet room, where they completed three tasks from the CogState task library. All three tasks featured virtual playing cards that were displayed sequentially on a computer monitor. Subjects were to provide ‘yes’ or ‘no’ responses to a task-dependent question using the right and left computer mouse buttons, respectively. The first task, Detection (DET), assesses sensorimotor processing speed using a simple reaction time test: subjects were to press only the ‘yes’ button as soon as possible after a card was displayed. The second task, Identification (IDN), assesses attention via choice reaction time: subjects were to indicate as soon as possible after each card presentation whether or not the card was red by pressing the appropriate mouse button. The last task, One-Back (ONB), additionally assesses declarative working memory: subjects were to press the appropriate mouse button to indicate whether or not the previous card was the same as the current card. The CogState software reports the mean button press accuracy and reaction time for each task. CogState also provides so-called “embedded invalidity indicators” (EIIs) which require minimum levels of accuracy and speed: DET and IDN require response accuracy to be greater than 90% and 80%, respectively. Mean reaction times in IDN and ONB must be greater than those in DET. We did not exclude any subject from further analysis if they failed to pass any CogState integrity check.

#### Robotic Assessment of Sensorimotor Learning

Subjects sat in a high-backed chair and grasped the handle of a horizontal planar robot, which they moved with their right hand (Fig 1A). An opaque screen mounted just above the plane of movement occluded direct view of both their hand and the robot arm. A computer projected visual stimuli onto the screen; these included a home target, a goal target, and (occasionally) a small cursor that provided real-time visual feedback of hand position. The home position and the target position were indicated using 0.4 cm diameter white dots spaced 10 cm apart along the line intersecting the horizontal and sagittal planes passing through the shoulder center of rotation. The goal target was positioned farther from the subject on that line than the home target. When displayed, the hand’s cursor appeared as a small white 0.4 cm diameter dot that accurately represented hand position.

**Fig. 1.**
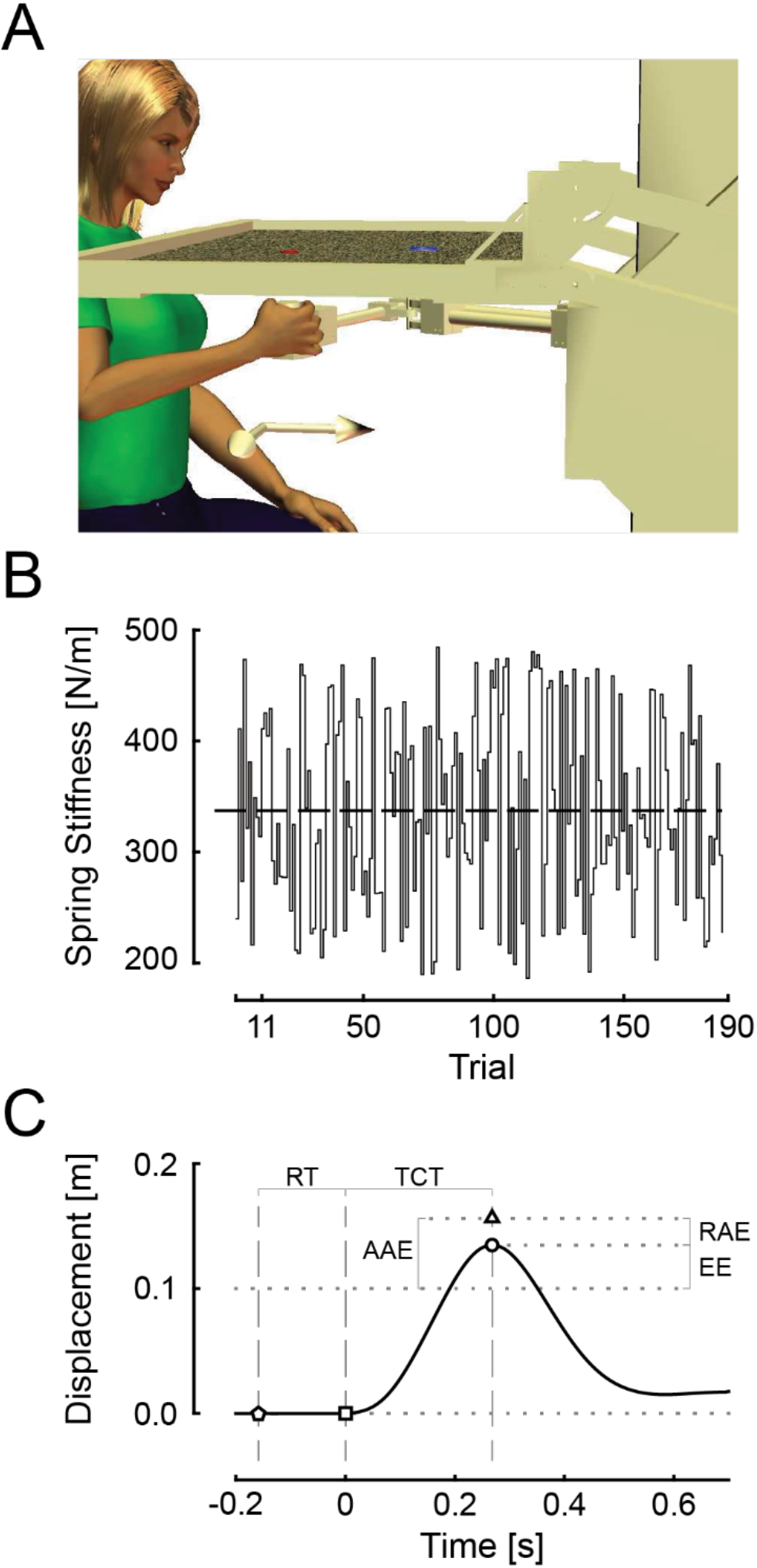
**A** Experimental setup: Home and goal targets were projected on a horizontal screen which occluded the subject’s view of the hand. A scintillating field was also projected to mitigate the presence of subtle, extraneous, visual landmarks on the screen. **B** Spring stiffness magnitude series used by the robot. Horizontal line indicates mean field strength of 338 N/m. **C** Hand displacement of a typical reach trial. Circle – point of maximum reach extent; triangle – point of self-assessed reach extent; square – point of movement onset defined as 10% of peak velocity; pentagon – point when the GO cue was presented. RT – reaction time from GO cue to movement onset; TCT – target capture time from movement onset to peak extent; EE – reach error relative to the target; AAE – absolute assessment error relative to the target; RAE – relative assessment error relative to the point of maximum extent.

We adapted the approach of Lantagne et al. (2021) to quantify the effects of intentional performance suppression on how implicit and explicit memories of kinematic performance contribute to sensorimotor adaptation. Specifically, we asked each subject to perform 190 experimental “trials” that were comprised of two phases each: a goal-directed out-and-back reaching movement followed by a self-assessment of kinematic performance. At the start of the reaching phase of each trial, subjects were to move the robot’s handle to the home target. Real-time visual feedback of hand position was provided by the cursor when the hand was within 2 cm of the home target; this was done to promote consistency of initial conditions across trials. Subjects were to hold at the home target and wait for a visual GO cue (the coincident disappearance of the home and goal targets), which signaled them to initiate a reach to the remembered goal location. Subjects were to move out and back to the target in one fluid movement to capture the goal target. Additional instructions differed between the two sessions (see Appendix A).

During each reach, the robot applied a spring-like resistance that opposed any movement away from the home target. The robot changed its rendered stiffness pseudorandomly between trials in a way that was unpredictable to the subject (Fig 1B). The trial sequence of stiffness values was drawn from a uniform distribution with a mean strength of 338 ± 86 N/m. All subjects experienced the same series of spring-like loads. The robot applied a virtual force channel (c.f. Scheidt et al. 2000) to constrain all movements of the handle to the straight line between the home and goal targets. This aspect of the experimental design constrained kinematic performance errors to those of extent, not direction, thereby simplifying subsequent analysis (cf. Judkins and Scheidt 2014). The robot recorded hand position and applied hand forces at a rate of 1000 samples per second. Immediately after the out-and-back reach phase, subjects were presented with graphical feedback of hand speed for two seconds, which informed them as to whether movements were made too quickly, too slowly, or within a specific desired range of speeds (0.8 m/s to 1.1 m/s). The purpose of this feedback was to promote movement consistency across the entire set of trials and across subjects.

After hand speed feedback was removed, subjects performed the self-assessment phase of the trial. Subjects were to use the robot’s handle to point to the peak extent of the prior reach, which they registered using a push-button response box held in their left hand. This procedure allows us to assess the accuracy of explicit recall of reach performance on each trial. The spring-like forces were removed during this phase so that only spatial information of the reach would be recalled, and not proprioceptive memory of the force experienced by the hand at peak extent.

During the first 10 reaching trials, the hand’s cursor was projected onto the screen in real time throughout the entire out-and-back movement. These “practice trials” were used to acclimate the subject to the spatial accuracy requirements of the task. For the remaining 180 trials (referred to as *test trials*), the cursor was removed coincident with delivery of the GO cue during out-and-back reaching. As described below, we analyzed performance in the test trials to assess the contributions of implicit memories to sensorimotor adaptation using the approach described in Lantagne et al. (2021).

### Data Analysis

#### Computerized Cognitive Assessment

We used the CogState Research2 toolkit to extract normalized measures of response time and performance accuracy from each of the three tests (i.e., DET, IDN, and ONB) performed in each session by each subject. The toolkit corrects for non-normality in the raw data by applying a log_10_-transform to response times and an arcsine-transform to the square root of the proportion of correct trials to total trials in each test (Maruff et al. 2009; Louey et al. 2014; Cromer et al. 2015). We used planned paired t-tests to compare the transformed performance measures across sessions to assess the effect of intentional performance suppression on CogState measures of sensorimotor processing speed (DET), choice reaction time (IDN), and declarative working memory (ONB).

#### Robotic Assessment of Sensorimotor Learning

Prior to post-processing, we used a semi-automated algorithm to identify individual trials wherein subjects failed to perform the task as instructed. We verified the algorithm’s recommendations through direct visualization of the hand’s displacement, velocity, and acceleration profiles in each reaching trial. Trials were excluded from further analysis for any of the following reasons: if the hand drifted more than 1 cm from the home position before the GO cue; if the acceleration profile of the out-and-back reach was not triphasic (i.e., suggesting corrective movements or a pause mid-reach); or if the subject pushed excessively against the channel constraints, causing the robot’s motor torques to exceed predefined safety limits (35 Nm). Across subjects, the exclusion criteria resulted in rejection of only 2.3% ± 3.9% of trials in the Best Effort session and 7.5% ± 8.4% of trials in the Performance Suppression session.

We extracted five kinematic outcome variables from the measured hand displacement trace in each trial. We defined *movement onset* as the moment where hand speed exceeded 10% of its first peak value (Fig 1C: square). *Reaction time* was defined as the time between the GO cue (Fig 1C: pentagon) and movement onset (Fig 1C: RT). *Target capture time* was the difference between movement onset and the moment the hand reached the maximum extent of outward movement (Fig 1C: circle, TCT). We quantified movement accuracy using *reach extent error* (*ε_i_*), defined as the signed difference between maximum movement extent and the actual target distance of 10 cm (Fig 1C: EE). Finally, we quantified *reach precision* as the standard deviation of the reach errors across trials within each session.

We additionally defined four kinematic parameters associated with the self-assessment of movement extent (Fig 1C: triangle). We quantified *absolute assessment error* (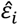; Fig 1C, AAE) as the signed difference between the recalled position of maximum reach extent (Fig 1C, triangle) and the target. We quantified the accuracy of subjects’ self-reporting using the *relative assessment error* (Fig 1C, RAE), which we defined as the signed difference between the self-assessed movement extent and the actual reach extent. We furthermore quantified the precision of these measures as the standard deviation of the absolute and the relative assessment errors, across trials within each session.

Subjects were excluded from further analysis if they met any of the following criteria: 1) all CogState measures improved when instructed to suppress performance; or 2) reach precision, target capture time, and reaction time all improved when instructed to suppress performance. Only one subject was excluded based on these criteria; that subject also self-reported after their Performance Suppression session that he had hardly attempted to emulate any of the symptoms he was asked to simulate (i.e., no symptom emulation effort was scored greater than 3 out of 10 for this subject).

### Models of sensorimotor adaptation and performance suppression

We examined the extent to which intentional performance suppression alters the way individuals process sensorimotor memories of reach performance as they adapt to changing environmental loads. We used systems identification techniques to compare the ability of four models of sensorimotor adaptation to capture the trial-by-trial variations in each individual’s performance in each robotic testing session (Eqn 1). Each model (Eqns 2a through 2d; Fig 2) estimates the relative contributions of sensorimotor memories to target capture performance during repetitive reaching under certain assumptions. They do so by fitting a unique function of kinematic memory 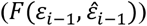 to the trial-by-trial performance data in each session:

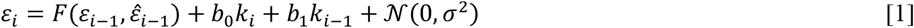

**Fig. 2.**
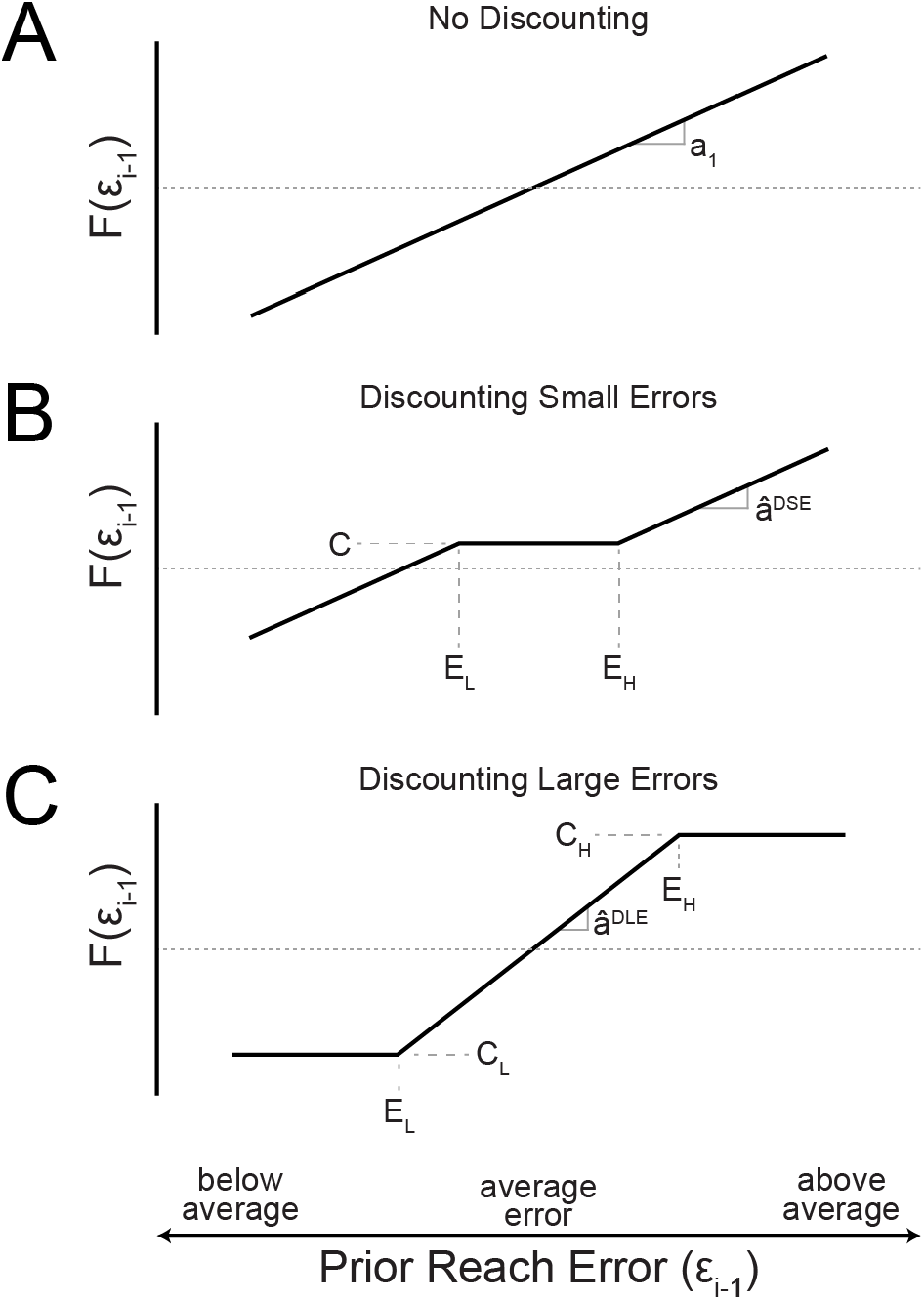
Linear and nonlinear memory-based models of sensorimotor adaptation during reaching. **A** Linear model (Eqn 2a and 2b); **B** discounting of small errors (Eqn 2c) within the range E_L_ to E_H_ with output bias, C, and feedback sensitivity of large errors, 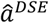; **C** discounting of large errors (Eqn 2d) outside the range E_L_ to E_H_ with corresponding discounted (clamped) output, C_L_ and C_H_, and feedback sensitivity for small errors, 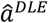.

Here, subscript *i* is a trial index spanning the range of test trials (i.e., from *i* = 11 to 190) such that *ε*_*i*–1_ and 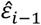 correspond to estimates of implicit and explicit memories of reach extent error from the previous trial, respectively. In all cases, we account for the impact of the robot’s physical resistance on movement by including input terms reflecting the current trial’s spring-like load (*k_i_*), as well as a memory of the robot’s load on the previous trial (*k*_*i*–1_). The residuals of the model fitting process were considered to be drawn from a normal distribution 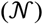 with zero mean and an observed variance *σ*^2^. Variables 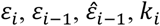, and *k*_*i*–1_ were centered by subtracting their respective means prior to model fitting.

The memory model of Equation 2a is the simplest model, describing a linear relationship between the objective kinematic performance error on any given trial *ε_i_* and the objective error on the prior trial (*ε*_*i*–1_) (Fig 2A; c.f., Scheidt et al. 2001). We regard the observed reach errors on the previous trial (*ε*_*i*–1_) as a proxy for implicit memory of reach performance (Lantagne et al. 2021). This model (i.e., Eqn 1 as augmented by Eqn 2a) has three parameters (*a*_1_, *b*_1_, and *b*_0_).

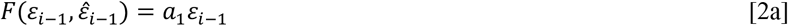

The model of Equation 2b additionally considers potential contributions from an explicit memory of reach performance, i.e., the absolute error of the self-assessed reach extent 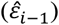 (Lantagne et al. 2021). As such, this model has four parameters (*a*_1_, *b*_1_, *c*_1_, and *b*_0_). Equations 2a and 2b both assess the possibility that intentional performance suppression could simply re-weight the contributions of explicit and/or implicit memories to subsequent performance.

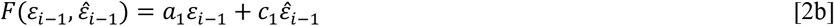

Equations 2c and 2d are non-linear models that consider the possibility of error discounting for either small errors (Eqn 2c and Fig 2B) or large errors (Eqn 2d and Fig 2C). Equation 2c instantiates a “satisficing” model (Goodrich et al. 1998) by implementing a dead band within which small errors within the range E_*L*_ ≤ *ε*_*i*–1_ ≤ *E_H_* are considered “good enough” and do not elicit proportional adjustments on the subsequent trial. The parameter ***C*** allows for a potential performance bias within the discounted range. The terms in parentheses implement the discontinuities between the discounted region and the error-sensitive regions on either side of the dead band. This model, which discounts small errors (DSE), has seven parameters (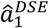, *E_L_, E_H_*, *C*, *b*_1_, *c*_1_, and *b*_0_).

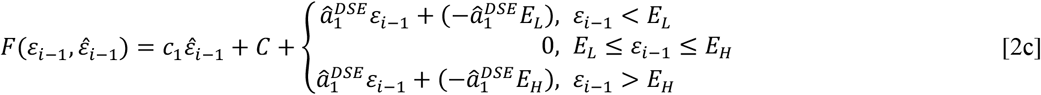

By contrast, Equation 2d yields a model that discounts large errors (DLE) outside the range E_*L*_ ≤ *ε*_*i*–1_ ≤ *E_H_* and is proportional within that range [Fig 2C; (c.f., Körding and Wolpert 2004)]. This model discounts large errors as fluke events that should not heavily influence subsequent motor movements. Here, C_*L*_ and C_*H*_ are constants representing the discounted responses beyond the sensitive region. The term in parenthesis implements the discontinuity between the sensitive and discounted regions (Fig 2C). The slope of the sensitive region for small errors, 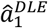, is defined as 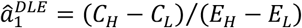 and is not an independent term in the model. Thus, this model also has seven parameters (*E_L_, E_H_, C_L_, C_H_*, *b*_1_, *c*_1_, and *b*_0_).

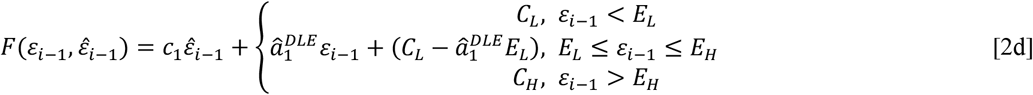

In each of the models, the personalized coefficient *b*_0_ relates how changes in the robot’s spring-like load contribute to changes in reach error on any given trial. Coefficients *a*_1_ (Eqns 2a and 2b), 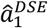 (Eqn 2c), and 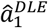 (Eqn 2d) describe the extent to which implicit memories of prior reach performance influence subsequent performance (cf., Lantagne et al., 2021). Coefficient *b*_1_ describes the extent to which memory of the prior robotic load impacts subsequent performance. Coefficient *c*_1_ describes the extent to which explicit memory of reach error on the prior trial impacts subsequent reach error.

We evaluated how well each model explained how subjects used memories of prior performance to compensate for environmental changes by computing the data variance accounted for (VAF) by model predictions (Eqn 3) for each session and each subject. Higher VAFs indicate better model performance.

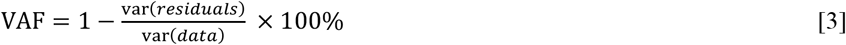

To facilitate comparison across models having different numbers of parameters, we adjusted the VAF for each model using a Minimum Descriptor Length (MDL) factor, which penalizes models having more parameters, thereby reducing the likelihood of model overfitting (Ljung 1999).

### Statistical Hypothesis Testing

We used planned, one-tailed pairwise t-tests to compare the normalized CogState measures of test timing and accuracy across the Best Effort and Performance Suppression sessions to determine whether subjects suppressed performance in each task as instructed. We similarly used planned pairwise t-tests to compare reach kinematic outcome measures across the two testing sessions to determine whether subjects suppressed performance in the reaching task. We used a repeated measures mixed effects ANOVA to compare subjects’ MDL-adjusted VAF across the four sensorimotor adaptation models to determine which model best described the observed data in the reaching task. We then fit the selected model to the data collected in each testing session for each subject to obtain personalized model coefficients for each testing session. We used two-tailed pairwise t-tests on each model parameter to test whether or not intentional performance suppression altered how sensorimotor memories contribute to the trial-by-trial updating of motor performance. All data processing, model fitting, and statistical processing was performed using MATLAB 2019a (The MathWorks, Natick, Massachusetts). Non-linear model fitting was performed with MATLAB’s *fminsearch* function minimizing the sum of squared errors. Statistical significance was set to a family-wise error rate of α = 0.05.

## Results

All subjects were attentive throughout experimental sessions and completed all required tests. After testing was completed for the Performance Suppression session, subjects most commonly reported that they attempted to suppress performance in that session by simulating fatigue, problems with memory, difficulty concentrating, and slowed responses (Fig 3). Actual performance on the CogState and robotic reaching tests exhibited the anticipated effects of this intentional performance suppression.

**Fig. 3.**
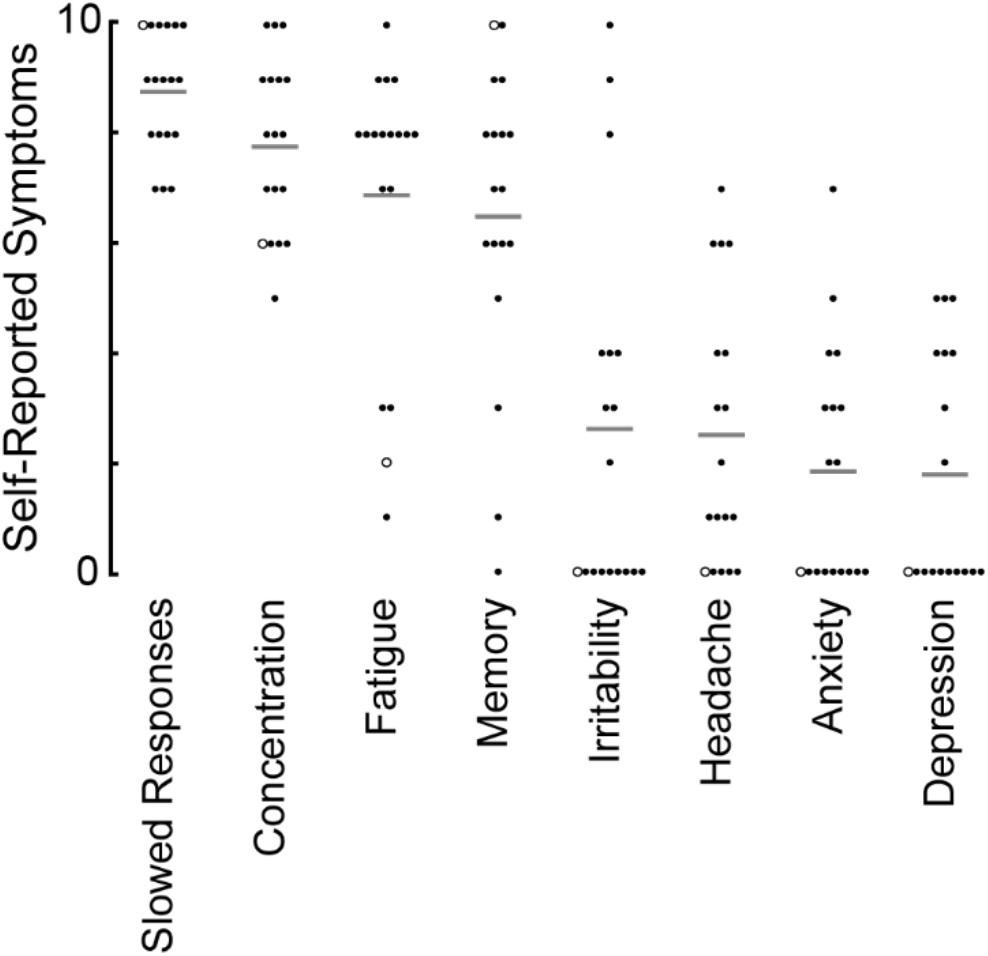
Cohort results of post-suppression-session survey of simulated symptoms (sorted from greatest effort to least effort). Grey lines: cohort-average score. Hollow circles: data for the selected subject shown in Figure 5.

### CogState scores

When instructed to perform with their best effort, all subjects passed each of CogState’s embedded invalidity indicators (EIIs). Best Effort scores in all three tests were comparable to those reported in previous studies using CogState to assess healthy individuals (Cromer et al. 2015) (Table 1). When instructed to intentionally suppress performance, subjects exhibited significant longer reaction times and commensurate less accurate responses relative to their best effort performances (all p < 0.001; Table 1; Fig 4). This was true not only for the subject cohort overall, but for the individual subjects with few exceptions [Fig 4: compare the prevalence of solid vs. dashed lines, which correspond to individual subject performance trends that were (or were not) consistent with performance suppression]. When suppressing performance, 12 of 18 subjects were flagged by at least one of CogState’s embedded invalidity indicators. The most commonly-flagged indicators were the Detection task’s 90% minimum accuracy requirement (9 subjects) and the reaction time comparison between the Identification and Detection task (7 subjects). We performed a follow-on analysis of performance trends in the six subjects who avoided triggering the CogState embedded invalidity indicators, finding that this subgroup significantly suppressed performance with respect to timing measures on all three tests but not with respect to the accuracy measures (speed: all p < 0.009; accuracy: all p > 0.065; Table 2).

**Table 1.** CogState performance of Best Effort and Suppression sessions across all subjects

| CogState Task | Measure | Best Effort | Suppressed | df | T Statistic | *p* |
| --- | --- | --- | --- | --- | --- | --- |
| DET | Speed | 2.49 (0.07) | 2.84 (0.18) | 17 | -7.605 | < 0.001 * |
|  | Accuracy | 1.51 (0.10) | 1.26 (0.26) | 17 | 4.087 | < 0.001 * |
| IDN | Speed | 2.67 (0.06) | 2.93 (0.16) | 17 | -5.944 | < 0.001 * |
|  | Accuracy | 1.46 (0.11) | 1.18 (0.23) | 17 | 4.256 | < 0.001 * |
| ONB | Speed | 2.80 (0.07) | 3.03 (0.17) | 17 | -4.777 | < 0.001 * |
|  | Accuracy | 1.33 (0.11) | 1.12 (0.18) | 17 | 4.596 | < 0.001 * |

Values reported as mean (SD). Table reports scores across all subjects regardless of EII flags (N = 18). CogState tasks: Detection (DET), Identification (IDN), and One Back (ONB). Better performance indicated by smaller speed and larger accuracy scores. P-value computed using one-tailed pairwise t-tests as Best Effort - Suppressed; \* indicates significant difference for $\alpha = 0.05$ ; left-tailed tests for Speed and right-tailed tests for Accuracy.

**Fig. 4.**
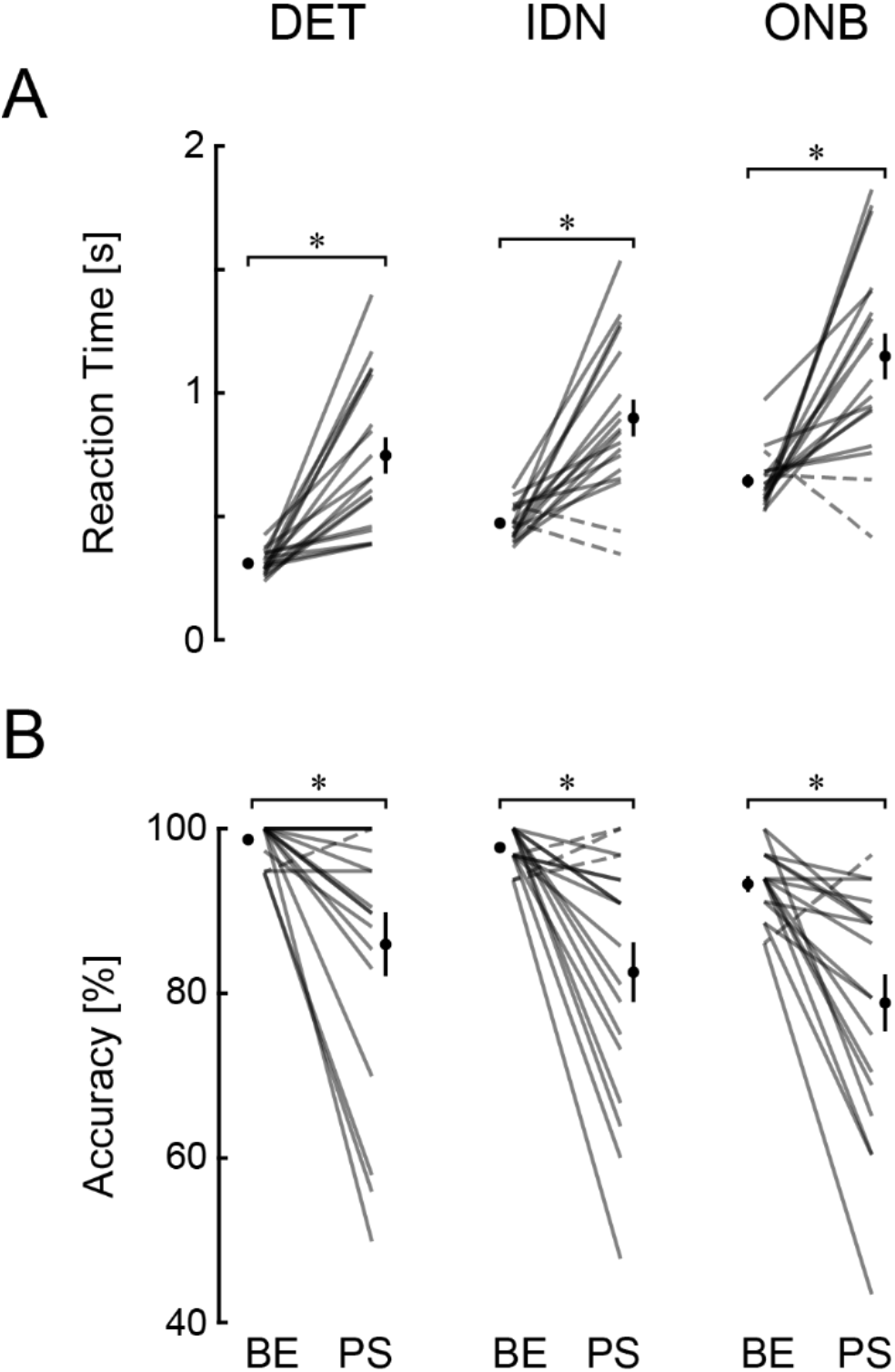
Cohort CogState raw timing (row **A**) and accuracy (row **B**) outcome measures for each test (column). DET: Detection; IDN: Identification; ONB: One-Back. Cohort means shown as solid circles. Error bars: ± 1 SEM. Note that some error bars fit within the circle used to denote the mean. Semi-transparent lines: individual subject performance. Solid lines: performance consistent with instructions to suppress performance; dashed line: performance that was inconsistent with instructions to suppress performance.

**Table 2.** CogState performance of Best Effort and Suppression sessions of subjects passing EIIs

| CogState Task | Measure | Best Effort | Suppressed | df | T Statistic | *p* |
| --- | --- | --- | --- | --- | --- | --- |
| DET | Speed | 2.45 (0.05) | 2.77 (0.09) | 5 | -6.031 | < 0.001 * |
|  | Accuracy | 1.53 (0.09) | 1.54 (0.07) | 5 | -0.214 | 0.840 |
| IDN | Speed | 2.65 (0.05) | 2.88 (0.13) | 5 | -3.484 | 0.009 * |
|  | Accuracy | 1.48 (0.10) | 1.32 (0.15) | 5 | 1.813 | 0.065 |
| ONB | Speed | 2.78 (0.04) | 3.03 (0.15) | 5 | -3.568 | 0.008 * |
|  | Accuracy | 1.34 (0.13) | 1.24 (0.10) | 5 | 1.474 | 0.100 |

Values reported as mean (SD). Table reports scores across subjects that did not fail an EII flag (N = 6). CogState tasks: Detection (DET), Identification (IDN), and One Back (ONB). Better performance indicated by smaller speed and larger accuracy scores. P-value computed using one-tailed pairwise t-tests as Best Effort - Suppressed; \* indicates significant difference for $\alpha = 0.05$ ; left-tailed tests for Speed and right-tailed tests for Accuracy.

### Robotic Reach Testing: Kinematic Performance

Kinematic data from a selected subject is shown in Figure 5 (panels A and B). Best Effort performance (reaction times: 412 ± 78 ms; target capture times: 273 ± 17 ms) was consistent with the generation of ballistic, out-and-back, target capture movements, which discourage feedback corrections during movement and instead foster corrective updating of motor plans between trials. The subject did, however, consistently overshoot the target spatially (average reach error: 6.1 ± 1.0 cm). This result is consistent with previous studies wherein visual feedback was withheld during reaching against unpredictable spring-like loads (Judkins and Scheidt 2014; Lantagne et al. 2021). When instructed to suppress performance, this subject had larger average reaction times (499 ± 170 ms) and target capture times (363 ± 56 ms). Note also that the variability of these measures was also larger relative to the Best Effort session. Although target capture accuracy was modestly smaller (average reach error increased by 11% to 6.8 cm in the Performance Suppression session), the spatial variability of peak hand displacement (± 1.9 cm) was larger in the Performance Suppression session. These outcomes are in accord with this subject’s self-reporting of how she intentionally suppressed performance (see hollow circle in Fig 3). Larger performance variability in the Performance Suppression session translated into larger trial-to-trial variability in the relationship between reach error and the robotic spring strength (Figs 5C and 5D).

**Fig. 5.**
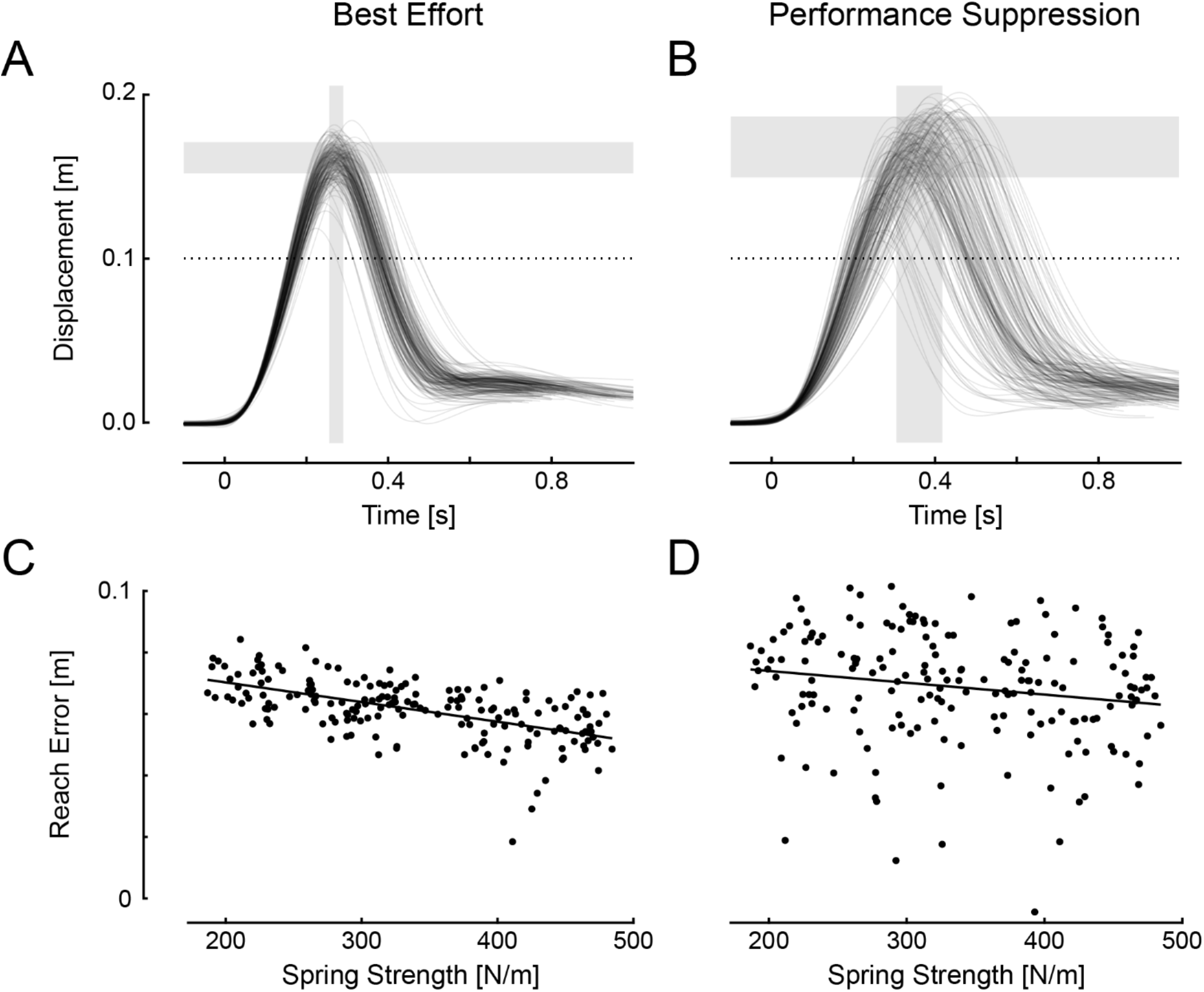
Measures of reach kinematics for a selected subject. **A-B**: Reach trajectories for Best Effort and Performance Suppression sessions, respectively. Visual feedback was not provided during reaching. For all reach traces, t = 0 seconds at movement onset. Horizontal dashed line at 10 cm indicates the target location. Horizontal grey bars indicate one standard deviation above and below the mean reach error. Vertical grey bars indicate one standard deviation before and after the mean target capture time. **C-D**: Reach error (peak reach extent – target distance) on each trial as a function of robotic spring stiffness for the Best Effort and Performance Suppression sessions. Black lines: best fit linear regression. Note the tendency to overshoot the target (bias) in both sessions.

The patterns of behavior displayed in Figure 5 were characteristic of those in the entire study cohort. Relative to Best Effort performances, intentional performance suppression resulted in longer reaction times (BE: 499 ± 170 ms; PS: 899 ± 527 ms; planned two-tailed paired t-test: T_17_ = 3.568, p = 0.001; Fig 6A), which were more variable (BE: 150 ± 85 ms; PS: 533 ± 586 ms; T_17_ = 3.143, p = 0.003; Fig 6F). Target capture times in the Performance Suppression session were also longer (BE: 295 ± 47 ms; PS: 382 ± 101 ms; T_17_ = 4.298, p < 0.001; Fig 6B) and had greater variability (BE: 29 ± 20 ms; PS: 67 ± 28 ms; T_17_ = 6.793, p < 0.001; Fig 6G) than Best Effort performances. This pattern of performance changes was observed for nearly every subject in our cohort (see individual-subject lines shown in Figs 6A, 6B, 6F, and 6G). Although we found no compelling evidence for a consistent change in mean reach error across sessions (BE: 6.3 ± 2.4 cm; PS: 6.0 ± 2.2 cm; T_17_ = 0.627, p = 0.539; Fig 6C; some subjects showed larger average reach errors in the Performance Suppression session whereas others had smaller average errors), the cohort had consistently larger reach error variability in the Performance Suppression session (BE: 1.4 ± 0.3 cm; PS: 1.9 ± 0.4 cm; T_17_ = 3.814, p < 0.001; Fig 6H).

**Fig. 6.**
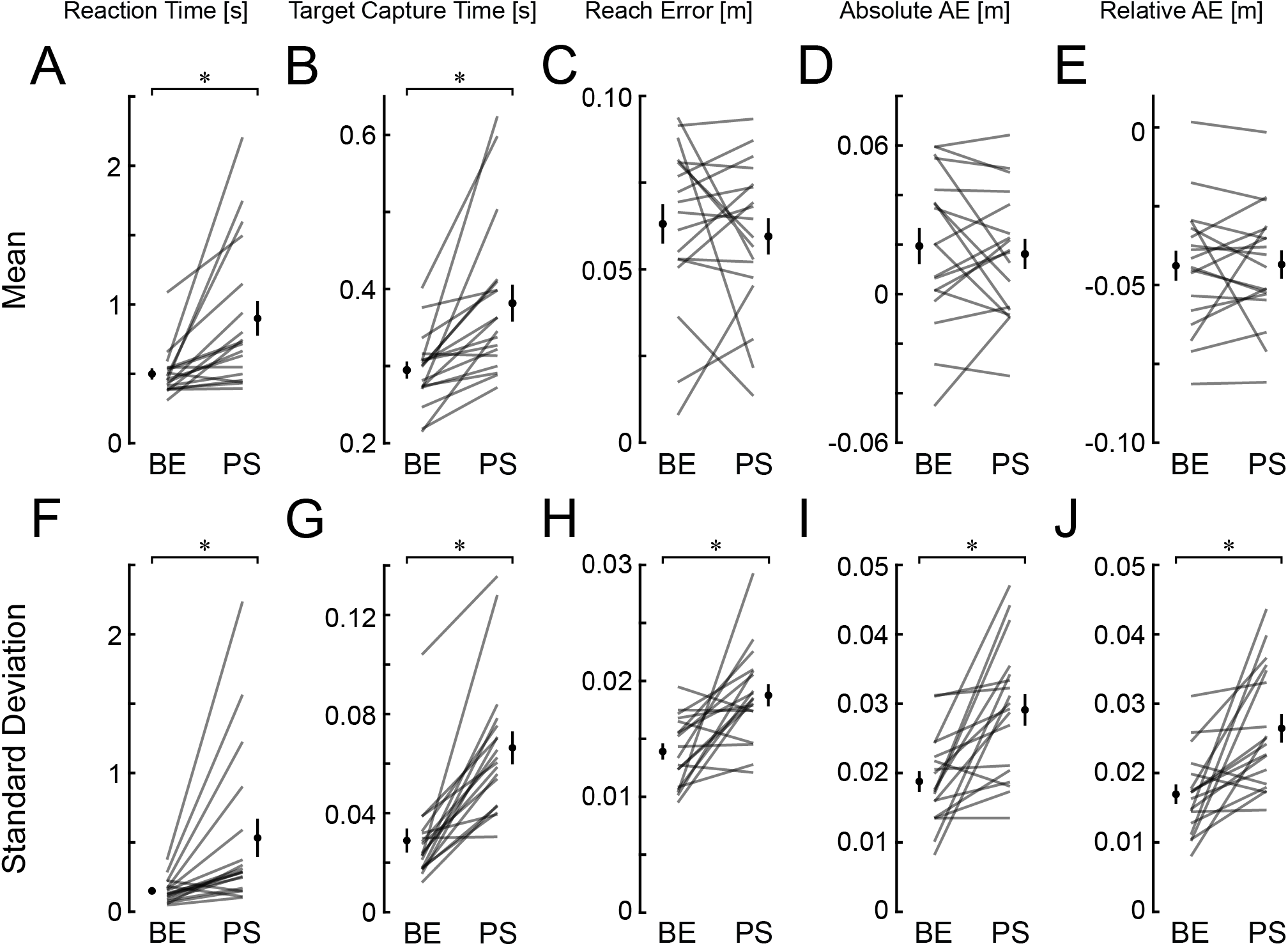
Cohort average measures of kinematic and self-assessment performance in the Best Effort (BE) and Performance Suppression (PS) sessions. Each column represents a kinematic or self-assessed measure with the top row showing the mean of the measure (accuracy) and the bottom row showing the standard deviation (precision). Solid circles: Cohort means. Error bars: ±1 SEM. Column 1: reaction time from the GO cue to movement onset. Column 2: target capture time from movement onset to moment of peak movement extent. Column 3: reach error at peak movement extent. Column 4: self-assessment of reach error (relative to the target). Column 5: relative assessment error between the reported self-assessment and actual reach error.

We similarly analyzed two measures of subjective performance derived from post-movement assessments of reach extent: absolute and relative assessment errors. Whereas absolute assessment errors provide an objective measure of explicit memory of prior reach extent, relative assessment errors account for trial-to-trial variations in actual reach extents that could be driven by trial-by-trial variations in the strength of robotic resistance to movement and/or in the vigor of executed movements. As noted for our objective measure of reach extent in Figure 6C, absolute assessment errors did not differ significantly between sessions (BE: 1.9 ± 3.1 cm; PS: 1.6 ± 2.6 cm; T_17_ = 0.601, p = 0.722; Fig 6D) although variability in this measure was larger when subjects were instructed to suppress performance (BE: 1.9 ± 0.6 cm; PS: 2.9 ± 1.0 cm; T_17_ = 4.274, p < 0.001; Fig 6I). Consistent with a prior study that interposed self-assessments of reach extent between trials (Lantagne et al. 2021), self-assessments in the current study markedly underestimated actual reach extents. The magnitude of this bias was about the same in both sessions: (BE: −4.4 ± 2.0 cm; PS: −4.3 ± 1.9 cm; T_17_ = 0.105, p = 0.459; Fig 6E). By contrast, the variability of relative assessment errors increased when subjects were instructed to suppress performance (BE: 1.7 ± 0.6 cm; PS: 2.6 ± 0.9 cm; T_17_ = 4.007, p < 0.001; Fig 6J). It is also worth noting that when we regress absolute self-assessment against prior reach extent error (Figs 7A), subjects bias the report of their reach extent as being more consistent than their objective performance (slope BE: 0.7 ± 0.4 cm; PS: 0.7 ± 0.2 cm; Fig 7B). They also report their reach extent closer to the target (intercept BE: −2.7 ± 3.7 cm; PS: −2.7 ± 2.2 cm; Fig 7C). These findings confirm those previously reported by Lantagne et al. (2021).

**Fig. 7.**
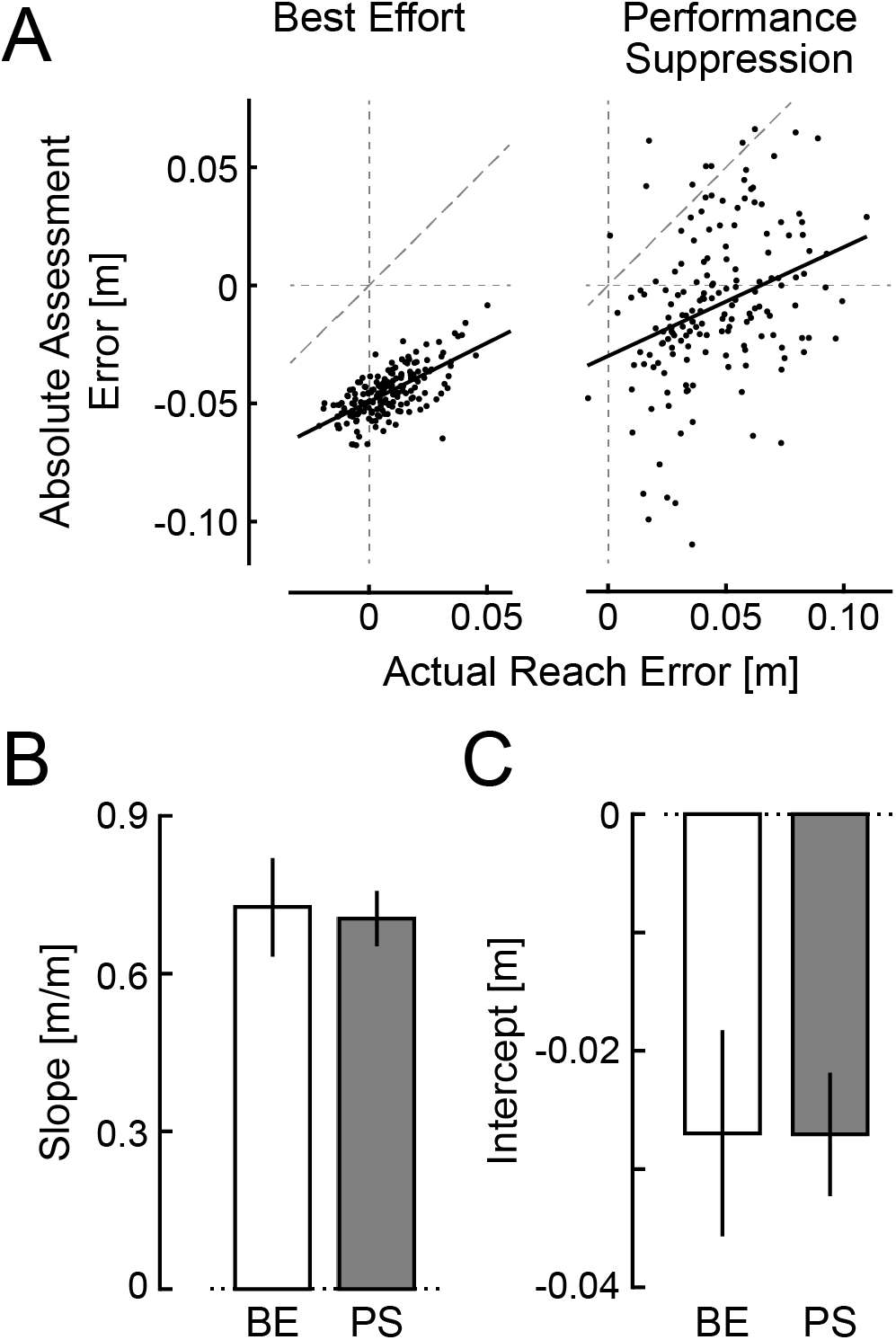
**A** Regression of self-assessment error onto reach error for a representative subject’s Best Effort session (BE; left) and Performance Suppression session (PS; right). Black solid lines: best fit linear regression. A dashed diagonal line with unity slope (perfect assessments of reach error) is shown for comparison. **B** Cohort results: slope of the relationship between self-assessed and actual reach errors between BE and PS sessions. **C** Cohort results: mean offset in the relationship between assessments and reach errors between sessions.

### Robotic Reach Testing: Models of Sensorimotor Adaptation

We next explored the extent to which intentional performance suppression might impact how implicit and explicit memories contribute to sensorimotor adaptation in response to variations in environmental resistance to movement during reaching. To do so, we compared the ability of four different memory-based models of sensorimotor adaptation (Eqns 1 and 2; see Fig 2) to account for variability inherent to the observed data (i.e., data variance accounted for; VAF) during both experimental sessions (Fig 8). Although two-way, repeated measures mixed-effects analysis provided evidence of a significant main effect of task instruction (i.e., experimental session) on model performance [F_(1,18.0)_ = 15.500, p < 0.001], it did not provide evidence for a systematic advantage for any of the higher-order models of Equations 2b – 2d, either as a main effect [F_(3,38.5)_ = 2.165, p = 0.108] or in interaction with session [F_(3,40.0)_ = 0.384, p = 0.765]. Notably, these results suggest that neither the inclusion of explicit memories of past reach error nor the inclusion of discounting strategies – either discounting for large or small errors – capture significant aspects of performance in our study. Consequently, we chose Equation 2a (Fig 8, diamond) as the most parsimonious model for use in analyzing the main effect of performance suppression on the contributions of sensorimotor memories to reach adaptation.

**Fig. 8.**
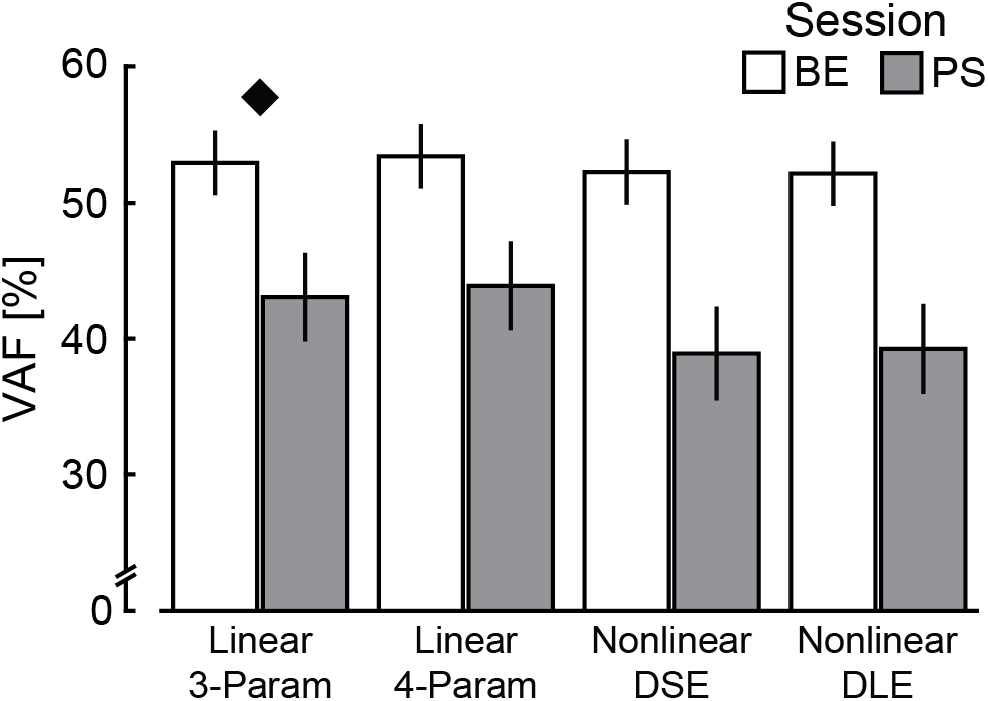
Variance accounted for (VAF) of each model in each session. Models in order from left to right correspond to the models of Equation 2. Diamond indicates the most parsimonious model selected for further analysis.

We then used multilinear regression techniques to fit the model of Equation 1 (per Eqn 2a) to each subject’s trial series of performance data. We included modulation terms (Eqn 4) to quantify the extent to which suppressing performance might change each model coefficient relative to the Best Effort testing session:

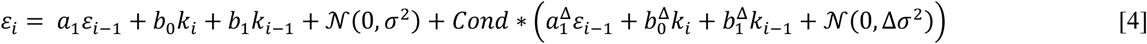

Here, *Cond* equals *0* for the Best Effort session and *1* for Performance Suppression. The multilinear regression yields eight parameters: The first four quantify coefficients *a*_1_, *b*_0_, *b*_1_, and *σ*^2^ when the subject performs with their best effort, whereas the last four quantify the *change* in model parameters (i.e., 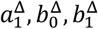 and Δ*σ*^2^) when the subject suppressed performance relative to their best effort. Figure 9 presents the results of this modeling for a selected subject, depicting observed reach errors and best-fit model predictions for both experimental sessions (Fig 9A), as well as the residuals from the fitting process (Fig 9B) and their corresponding frequencies (Fig 9C). Lilliefors test of normality was used to determine if each subject’s residuals followed a Gaussian distribution. We drew two conclusions from scrutiny of all the subjects’ results at this level: 1) the distribution of the residuals can be described as Gaussian for all subjects during the Best Effort session and Gaussian for the most subjects (13 of 18) during the Performance Suppression session, 2) the variance of residuals is consistently greater during the Performance Suppression session relative to the Best Effort session.

**Fig. 9.**
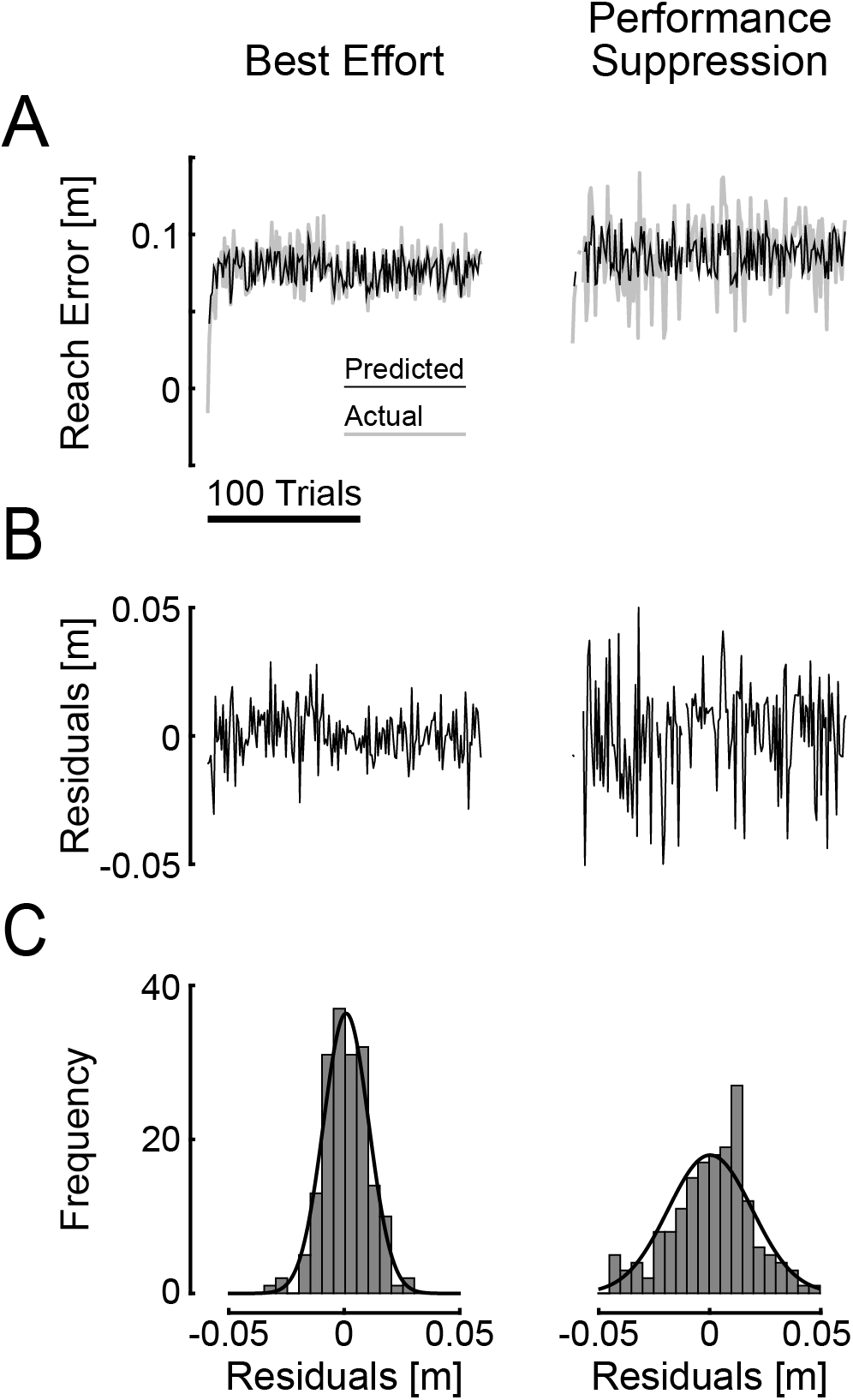
Time series analysis. A selected subject’s time series of reach errors, model predictions, and residuals of Equations 1 and 2a for Best Effort (left column) and Performance Suppression (right column). **A** Actual reach errors (gray lines) and model predictions (black lines). **B** Residuals of the model fit as a function of trial number. Trial series scale bar as displayed in panel A. **C** Distribution of the residuals with best-fit Gaussian function.

These conclusions were supported by analyses of personalized estimates of model parameters (Fig 10). Planned two-sided t-test found that all four BE coefficients (*a*_1_, *b*_0_, *b*_1_, and *σ*^2^) were significantly different from 0 (T_17_ > 11.00, p < 0.001 in each case) indicating that each parameter was needed to model the trial-by-trial variations in performance observed in our study. By contrast, the coefficients 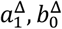, and 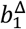 did not differ significantly from 0 (T_17_ < 1.83, p > 0.085 in each case), indicating efforts to suppress performance did not substantially change how the prior error, current spring, and prior spring, respectively, influenced subsequent performance. However, *Δσ*^2^ was significantly different from 0 (T_17_ = 4.82, p < 0.001). Parameter means and standard deviations are shown in Table 3. When trying to suppress performance, subjects did not apparently change how memories of prior reach performance influenced subsequent reach attempts, but instead increased variability of reach errors in a way that could not be accounted for by the systematic contribution of sensorimotor memories.

**Fig. 10.**
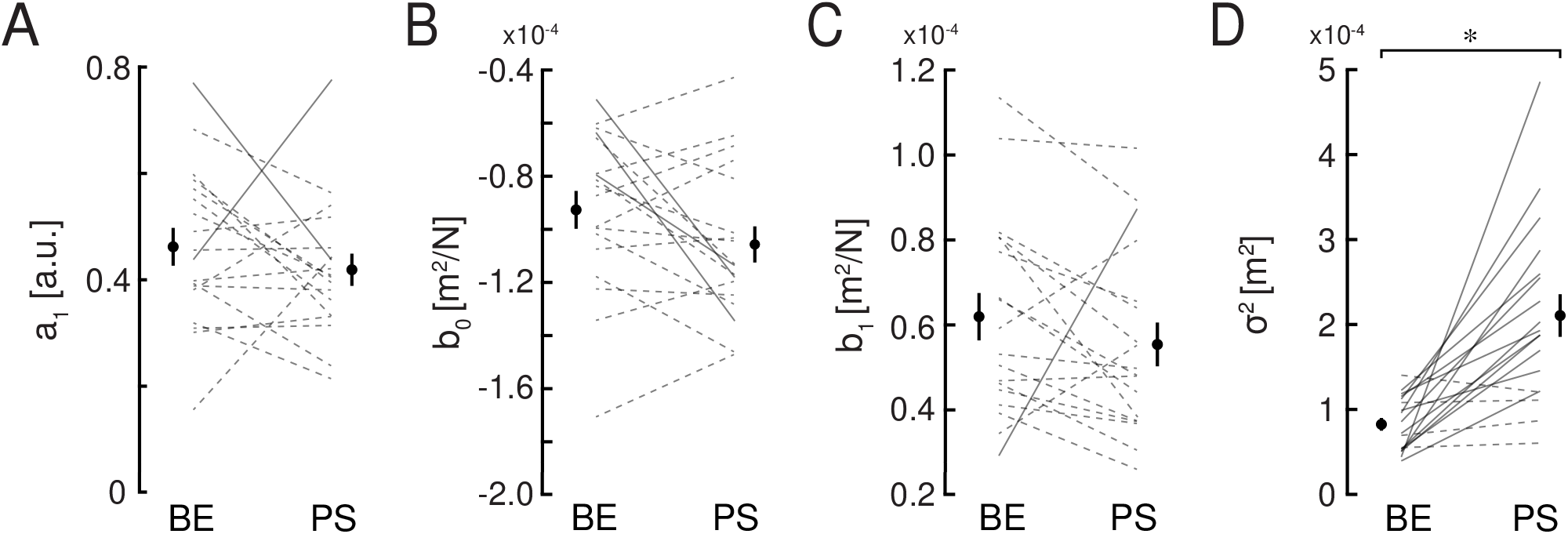
Cohort results: Adaptation model coefficients of each subject from Equation 4 compared for Best Effort (BE) and Performance Suppression (PS) sessions. Solid lines indicate a significant change of the coefficient between sessions after Bonferroni correction; dotted lines indicate no change. Solid circles: Cohort averages. Error bars: ±1 SEM. **A** model coefficient *a*_1_ – relative contribution of the previous reach error. **B** model coefficient *b*_0_ – relative contribution of the present spring stiffness. **C** model coefficient *b*_1_ – relative contribution of the previous spring stiffness. **D** model residuals within session.

**Table 3.** Contributions of sensorimotor memories compared across Best Effort and Suppression sessions

| Coefficient | Best Effort | Suppressed | df | T Statistic | *p* |
| --- | --- | --- | --- | --- | --- |
| $a_1$ (a.u.) | 0.462 (0.151) | 0.419 (0.129) | 17 | 1.036 | 0.315 |
| $a_1^\Delta$ (a.u.) | -0.043 (0.177) | | | | |
| $b_0$ (cm<sup>2</sup>/N) | -0.927 (0.303) | -1.057 (0.289) | 17 | 1.830 | 0.085 |
| $b_0^\Delta$ (cm<sup>2</sup>/N) | -0.130 (0.302) | | | | |
| $b_1$ (cm<sup>2</sup>/N) | 0.619 (0.237) | 0.553 (0.218) | 17 | 1.267 | 0.222 |
| $b_1^\Delta$ (cm<sup>2</sup>/N) | -0.066 (0.222) | | | | |
| $\sigma^2$ (mm<sup>2</sup>) | 83.0 (32.0) | 211.0 (106.5) | 17 | 4.823 | < 0.001 * |
| $\Delta\sigma^2$ (mm<sup>2</sup>) | 128.0 (112.6) | | | | |

Values reported as mean (SD). \* indicates significant difference for $\alpha = 0.05$ .

In about half of our subjects’ sessions, we noted a slight, slow drift in reach errors over the course of the 180 test trials. To determine the impact of this nonstationarity on the pattern of results describe above, we repeated the multilinear regression analyses after removing linear trends from the time series data. The results of these additional analyses yielded the same pattern of results as described above. Thus, slow drifts in performance do not significantly impact the conclusions drawn regarding the sensitivity of memory-based adaptation to intentional performance suppression.

### Validation Against Previously Published Findings

As a further validation, we compared model coefficients obtained during the Best Effort sessions to model coefficients obtained by Lantagne et al. (2021) under nearly identical “no visual feedback” conditions (their “No Visual Feedback, Proprioceptive Assessment” condition, NV-PA) and to coefficients obtained in a condition where subjects were provided with visual feedback of a cursor representing the hand’s true location (i.e., their “Visual Feedback, No Assessment condition, V-NA). For each coefficient (*a*_1_, *b*_0_, and *b*_1_), one-way ANOVA identified a significant difference between conditions {Best Effort, NV-PA, V-NA}: *a*_1_ [F_(2,55)_ = 19.801, p < 0.001]; *b*_0_ [F_(2, 55)_ = 3.284, p = 0.045]; and *b*_1_ [F_(2, 55)_ = 4.890, p = 0.011]. Post-hoc tests revealed these condition effects were driven entirely by the availability of visual feedback in the V-NA condition but not in the other two cases. Using the Best Effort condition as a reference for comparisons, only coefficients in the V-NA condition of Lantagne et al., 2021 differed significantly from the Best Effort coefficients reported here, whereas the task-similar NV-PA condition yielded comparable coefficient values: *a*_1_ (NV-PA: p = 0.173; V-PA: p < 0.001); *b*_0_ (NV-PA: p = 0.985; V-NA: p = 0.034); and *b*_1_ (NV-PA: p = 0.417; V-NA: p = 0.037). The results of this validation analysis support the conclusion that the effects of intentional performance suppression were minimal with regards to how implicit memories contribute to sensorimotor adaptation in our task, i.e., less than the effects of providing concurrent visual feedback of hand motion during repeated reaching against unpredictable spring-like loads.

## Discussion

We investigated the extent to which intentional performance suppression impacted how implicit and explicit memories of sensorimotor performance contribute to adaptation of reaching movements. Healthy human subjects participated in two testing sessions where they performed a series of computerized cognition tests and a robotic test of goal-directed reaching. The cognition tests assessed simple and choice reaction times and the integrity of short-term memory; they also helped to confirm that subjects understood and complied with task instructions. The robotic test assessed how implicit and explicit memories of sensorimotor performance contribute to the adaptation of goal-directed reaches to unpredictable changes in the magnitude of a hand-held load. In one session, subjects performed to the best of their ability. In the other session, subjects were to suppress performance by emulating common symptoms of mild traumatic brain injury (an approach adapted from Suhr and Gunstad 2000). Relative to their best effort, subjects typically exhibited slowed and less accurate responses in the computerized cognition tests, as well as slowed and more variable movements in the robotic test of sensorimotor adaptation. Subjects also self-reported emulation of symptoms including slowed responses, difficulty concentrating, fatigue, and memory problems. We used systems identification techniques (cf., Ljung, 1999) to compare the ability of four memory-based models of sensorimotor adaptation to capture the evolution of reach errors from one trial to the next. In both sessions, subjects primarily used implicit memories from the most recent trial to adapt their reaches to unpredictable changes in the robot’s resistance to motion; these results confirm and extend prior reports using similar best-effort approaches (Thoroughman and Shadmehr 2000; Scheidt et al. 2001; Judkins and Scheidt 2014; Lantagne et al. 2021). The modeling results showed that subjects did not “give up” or otherwise discount the presence of large errors in either session, nor did they use a “good enough” strategy that discounted small errors (i.e., satisficing, Goodrich et al. 1998). Including a term related to explicit memory also did not improve model performance over the simplest model that engaged only implicit memories (Eqns 1 and 2a). Importantly, and despite clear evidence of performance suppression in measures of response timing and movement variability, modeling revealed no systematic changes in how sensorimotor memories contribute to reach adaptation across the two sessions. Rather, each subject suppressed performance in a way that seemed to increase the variability of movement vigor randomly from one trial to the next. We validated our results against previously published data, further supporting the conclusion that intentional performance suppression had minimal impact on how implicit sensorimotor memories contribute to adaptation in our study.

### Contributions of explicit and implicit processes to the adaptation of reaching movements

Numerous previous studies have shown that distinct explicit and implicit processes can contribute to sensorimotor adaptation (Smith et al. 2006; Mazzoni and Krakauer 2006; Hwang et al. 2006; Mazzoni and Wexler 2009; Benson et al. 2011; Taylor et al. 2014; McDougle et al. 2015; see Maresch et al. 2021 for a review). In one of these, Taylor and colleagues (2014) used a technique capable to isolate the contributions of explicit and implicit processes to the adaptive compensation for a 45° visuomotor rotation during horizontal planar reaching. Subjects were instructed to make 7 cm reaches while holding a digitizing stylus in their right hand. Taylor and colleagues asked subjects to report - prior to each reach - where they intended to aim on the following movement using an angular reticule of visual landmarks that was projected onto a visual display placed just above the plane of hand movement. The display screen also provided cursor feedback of hand movement during and/or after some movements. Subjects were not informed about the rotation prior to its occurrence, nor were they given suggestions about any strategy to counter it. The authors found that upon imposing the rotation, subjects readily chose to aim to locations other than the goal target. The authors interpreted the intended aiming direction as a faithful representation of an explicit movement plan. The difference between the explicit aiming direction and the actual reach direction was assumed to be due to implicit mechanisms of sensorimotor adaptation. Over the course of 320 movement trials with imposed rotation, the magnitude of the verbalized aim angle rapidly increased then slowly decreased. By contrast, the contributions of implicit sensorimotor adaptation to the overall compensatory response increased only slowly over the series of movements such that explicit re-aiming dominated in the first half of the movements whereas implicit sensorimotor adaptation dominated in the later movements.

We adjusted the approach of Taylor et al. (2014) to have a retrospective focus, i.e., by asking subjects to recall and report the kinematic outcome of the previous movement rather than to indicate their plans for the upcoming movement. Our goal was to narrowly assay explicit memories of sensorimotor performance that could potentially contribute to any compensatory response, whether an explicit strategy of re-aiming or an implicit process of sensorimotor recalibration. We asked subjects to use their hand to point to the recalled location to obtain a precise measure of explicit memory without relying on imprecise verbalizations. Consistent with findings reported in a recent study using similar techniques (Lantagne et al. 2021), subjects in the current study recalled peak movement extents that were biased closer to the target than those that were actually performed. We used the objective (actual) reach error as a proxy for an implicit memory of reach error; remarkably, stepwise multilinear regression revealed that memories of prior reach errors were more predictive of trial-by-trial variations in the upcoming reaches than explicitly recalled errors. Further, the addition of explicit memories of past reach error provided no additional explanatory power to models of trial-by-trial adaptation, either when subjects were instructed to perform to the best of their ability or when they actively suppressed performance.

As demonstrated recently by Maresch and colleagues (Maresch et al. 2020), different experimental outcomes can be obtained when using of direct verbal reporting vs. indirect approaches to measuring the contributions of explicit re-aiming and implicit adaptation to the overall compensation for an environmental perturbation. It is likely that such methodological differences can explain why we found evidence only for implicit memory contributions to sensorimotor adaptation and the insensitivity of that adaptation to intentional performance suppression whereas Taylor et al (2014) report evidence for both implicit and explicit process contributions to adaptation. One major difference stems from the fact that load perturbations in our study were designed to impact only movement extent whereas the visuomotor rotation in the Taylor study primarily impacted movement direction. A growing body of experimental evidence suggests that movement direction and extent are planned and controlled separately (Bock and Arnold 1992; Gordon et al. 1994; Messier and Kalaska 1997; Bhat and Sanes 1998; Sainburg et al. 2003; Poh et al. 2017) and that they engage separate neural mechanisms (Krakauer et al. 2004; see also Desmurget et al. 2003; Schlerf et al. 2012). As such, it is possible that different adaptive mechanisms may have been recruited and assayed in the two studies.

Another difference between the current study and previous assessments of interactions between explicit and implicit adaptive processes (e.g., Mazzoni and Krakauer 2006; Keisler and Shadmehr 2010; Benson et al. 2011; Taylor et al. 2014) is that subjects in our study adapted to loads that changed unpredictably from one trial to the next whereas subjects in most past studies adapted to a predictable perturbation (such as a constant visuomotor rotation within the range 30° to 60°). When the load change is predictable, an explicit re-aiming strategy can rapidly reduce performance errors, although a slower process of implicit recalibration still occurs (Mazzoni and Krakauer 2006), albeit to a potentially lesser extent (Benson et al. 2011). The developmental time courses of these fast and slow compensatory processes were identified by Keisler and Shadmehr (2010), who also showed that only the faster compensatory process can be interfered with by engaging declarative memory in a secondary task (Keisler and Shadmehr 2010). In that study, subjects grasped the handle of a planar robot and performed outward reaches to a single target. The robot sometimes applied a viscous curl forcefield that acted to push the hand perpendicular to the direction of reaching. Subjects were first exposed to a predictable field A and they learned to compensate for the perturbation over the course of nearly 400 trials (see Figures 2 and 3 in Keisler and Shadmehr 2010). This was followed by a brief, 20-trial exposure to an opposite field B (B = -A), which drove the adaptive state of the fast process to oppose the state built up by the slow process, driving the overall compensatory response toward zero within approximately 12 trials (see also Smith et al. 2006). This set the stage for the main intervention of the study: over the course of the following 3 minutes, one group of subjects engaged declarative memory by performing a verbalized word-pair recall task; another group performed a nonmemory task that required them to count and report the number of vowels in a series of nonsense character strings; a control group simply rested for 3 minutes. Subjects then made a series of reaching movements in the presence of a mechanical error clamp (Scheidt et al. 2001; Smith et al. 2006) that allowed observation of the overall compensatory response without disturbing the adaptive states themselves. During the error clamp trials, the contributions of the fast adaptive process to the overall compensatory response were isolated from those of the slow component by measuring “spontaneous recovery”, whereby peak hand forces observed in the channel trials initially increase before slowly decreasing toward baseline. Only the declarative memory task interfered with the fast component contributions to the overall adaptive response; interference was not noted for either the nonmemory or the control groups. No interference was observed for the slow process in any group, demonstrating the distinction between memory resources serving explicit and implicit sensorimotor learning.

The use of random perturbations in our study likely discouraged attempts to form an explicit memory-based strategy to compensate for the seemingly random upcoming load (see also Lantagne et al. 2021). However, when instructed to actively suppress performance, subjects readily complied by making slowed and more variable movements. Our modeling results showed that subjects did not discount the presence of large or small errors in either session, although it is possible that the range of errors produced by our set of experimental perturbations (average range = 9.3 ± 2.8 cm) were contained within a very wide region of linear sensitivity and thus we did not have an opportunity to observe discounting of large errors. Including a term related to explicit memory also did not improve model performance over the simplest model that engaged only implicit memories of performance on the most recent trial to predict and compensate for the upcoming mechanical load (Eqns 1 and 2a; see also Scheidt et al. 2012). Indeed, it is hard to imagine that subjects could effectively reweight the systematic influence of sensorimotor memories (Eqns 1 and 2a) over the course of 100+ movements based on explicit recall of past performance when the perturbations change unpredictably from one trial to the next. The lack of a measurable difference in the relative contributions of implicit sensorimotor memories across the Best Effort and Performance Suppression sessions was not due to a lack of sensitivity in our approach. To test this, we compared model coefficients obtained during the Best Effort sessions here to coefficients obtained by Lantagne et al. (2021) under an identical “no visual feedback” condition as well as to a condition where subjects were provided with visual feedback of a cursor representing the hand’s true location. As in that previous study, we found that relative to testing conditions without a concurrent visual feedback of hand position, simply adding a cursor changed how subjects combine implicit memories of performance as they try to adapt to the unpredictable loads.

It is interesting to consider the time course over which the fast and slow adaptive processes evolve, as described in studies using deterministic perturbations (fast: ~10 trials to reach asymptote; slow: at least 30 to 40 trials to reach asymptote; see (Krakauer et al. 2000; Smith et al. 2006; Keisler and Shadmehr 2010; Taylor et al. 2014; McDougle et al. 2017; Yin and Wei 2020). By contrast, the implicit process described by Equations 1 and 2a asymptotes within 6 to 10 trials (see Scheidt et al. 2001, their Figure 7A), which is similar to that of the fast process described by Smith and colleagues (2006). It is not yet clear as to why the implicit form of adaptation tested here should be insensitive to intentional performance suppression while also developing with a time course similar to that described for the fast labile process described previously (e.g., Smith et al. 2006; Keisler and Shadmehr 2010), although as mentioned above, differences in experimental approach may contribute (e.g., a focus on movement extent rather than direction; stochastic vs. deterministic perturbations, retrospective vs. prospective reporting, etc.).

### Limitations and Future Directions

There are a number of limitations in our work. We assumed but did not test whether explicit reporting might impede the adaptive processes we sought to assay. We did so however in a prior study by comparing the influence of sensorimotor memories on adaptation across testing conditions that either did or did not require explicit reporting of reach extent; no significant differences were observed (Lantagne et al. 2021). A critical reader might also note that report-based measures are subject to bias (Metcalf et al. 2007; Hadjiosif and Krakauer 2021; Lantagne et al. 2021). However, our systems identification procedures subtract the mean from each regressor prior to modeling, thereby making them insensitive to bias in the behavioral time series, including the recalled and reported movement extents. A more substantial limitation is the unknown source of increased variability of model residuals when subjects suppressed their performances. Examination of the model residuals did not reveal systematic deviations from normality, suggesting that there was no systematic (yet unmodeled) source of the variability. Because experimental psychologists generally maintain that people cannot behave randomly, at least without specialized feedback of their performance (Neuringer 1986), further investigation is warranted to better understand how subjects may be able to increase reach variability in this task in a seemingly random way.

This work was motivated in part by the possibility that greater understanding of the interactions between explicit and implicit learning processes will have practical utility in the assessment of neurologic injury, for example after sport-related concussion (McCrea et al. 2004; Register-Mihalik et al. 2013). It has been reported that a significant proportion of athletes intentionally suppress pre-season baseline performance on computerized cognitive tests and underreport post-concussion symptoms so they can return to play as early as possible (Schatz and Glatts 2013; Higgins et al. 2017; Raab et al. 2020). There exists a need for an objective measure of neurocognitive function that is robust against explicit strategies of performance suppression. Future studies may wish to explore the extent to which an assay of sensorimotor adaptation of movement in the presence of unpredictable environmental perturbations can detect abnormalities due to neurologic injury and their resolution with recovery.

## Declarations

### Funding

This work was supported in part by a grant from the Marquette University Athletic and Human Performance Research Center (SIP105), and by the National Science Foundation (NSF) under an Individual Research and Development Plan. Any opinions, findings, conclusions, or recommendations expressed in this material are those of the authors and do not necessarily reflect the views of the NSF.

### Conflicts of interest/Competing interests

Access to the CogState Research2 toolkit was provided by CogState, Inc. without cost or obligation. The authors declare no conflicts or competing interests.

### Ethics approval

The study protocol was approved by the IRB at Marquette University (HR-3233).

### Consent to participate

All subjects gave informed written consent to participate in this study.

### Consent for publication

Each author has read and concurs with the content in the manuscript.

### Availability of data and material

De-identified data will be made available upon reasonable request.

### Code availability

N/A

## Acknowledgements

We thank the participants in this study and the Marquette University Psychology Department for assisting in subject recruitment.

## Authors’ contributions

Conceived of study: D Lantagne, L Mrotek, J Hoelzle, D Thomas, R Scheidt; robot programming: D Lantagne, R Scheidt; data collection: D Lantagne; analyze and interpret data: D Lantagne, L Mrotek, R Scheidt; wrote manuscript: D Lantagne, L Mrotek, J Hoelzle, D Thomas, R Scheidt; approved manuscript: D Lantagne, L Mrotek, J Hoelzle, D Thomas, R Scheidt.

## Appendix A

### Instructions for Best Effort Session

While you are participating in our study today, we ask that you try to do your very best.

<CogState> Perform these next computerized tests as quickly and accurately as you are able.

<Robotic Reach Testing> For this next test, move out and back to the target in one fluid movement to capture the goal as quickly and accurately as possible. After each movement, we will ask you to move the handle to indicate how far you just moved. Try to be as accurate as possible.

### Instructions for Performance Suppression Session

For today’s session, I want you to imagine that you were in a car accident in which another driver hit your car. You were knocked unconscious and woke up in the hospital. Doctors told you that you experienced a mild traumatic brain injury. You were kept overnight for observation.

You are now involved in a lawsuit against the driver of the other car. As a part of the lawsuit, you are required to undergo testing to determine the severity of your brain injury. You have decided to fake or exaggerate symptoms of a brain injury in an attempt to increase the settlement you will receive. If you can successfully emulate lingering effects of brain injury, you are likely to get a better settlement. If you are caught faking however, you are likely to lose the lawsuit.

<CogState> You are about to take a series of tests that can assess effects of mild traumatic brain injury. Common effects include: Slowed responses, difficulty concentrating, memory problems, fatigue, headache, irritability, anxiety, and depression. During the following tests, I would like you to emulate these effects in a believable way such that I cannot tell that you are faking.

<Robotic Reach Testing> For this next test, move out and back to the target in one fluid movement to capture the goal target. After each movement we will ask you to move the handle to indicate how far you just moved. Recall that common effects of mild traumatic brain injury include: Slowed responses, difficulty concentrating, memory problems, fatigue, headache, irritability, anxiety, and depression. During the following test, I would like you to emulate these effects in a believable way such that I cannot tell that you are faking.

## Notes

### Competing Interest Statement

The authors have declared no competing interest.

